# scCircaSim: A Circadian Simulator for Single-Cell and Spatial Transcriptomics

**DOI:** 10.64898/2026.11.05.764753

**Authors:** Chenyu Li, Liwei Zhang, Qian Zhang, Liang Zhao, Xiaoxia Song, Yu Tian, Guozhang Tang, Hong Zhou, Zhibin Wang, Pin Sun

## Abstract

**Motivation:** Simulated single-cell and spatial transcriptomic datasets that capture circadian dynamics while providing known ground truth remain scarce, limiting systematic evaluation of circadian analysis methods.

**Results:** We developed scCircaSim to generate circadian single-cell and spatial transcriptomic data from a single-time-point reference. The framework combines phase progression with reference-derived network propagation of anchor gene rhythms and incorporates neighborhood effects for spatial simulation. Outputs retain reference heterogeneity and provide known cell or spot phases, rhythmicity labels, and gene-level mesor, amplitude, and phase values. Validation showed retention of reference structure and recovery of specified temporal patterns. Benchmarks of four cell phase inference methods and three rhythmic gene analysis methods revealed differences in phase accuracy, rhythmicity detection, and parameter recovery. scCircaSim supports reproducible method evaluation and preliminary experimental planning.

**Availability and Implementation:** scCircaSim is implemented in R and available at https://github.com/Liwei-DynamicsLab/scCircaSim.

**Supplementary Information:** Supplementary figures accompany the manuscript.

## Introduction

Circadian rhythms coordinate mammalian physiology with the daily light and dark cycle(1). Through interactions between central and peripheral clocks, temporal regulation shapes transcription, metabolism, immune activity and tissue-specific functions(2, 3). Its disruption is associated with diverse pathological conditions, motivating investigation of how rhythmic programs vary among cells and spatial microenvironments(4, 5). Bulk RNA sequencing has enabled systematic identification of rhythmic genes(6), while single-cell and spatial transcriptomics increasingly reveal temporal heterogeneity beyond population averages(7, 8).

Computational analysis for circadian rhythmicity has developed along two complementary directions. CYCLOPS, TEMPO, Cyclum and CoPhaser reconstruct cellular phase using latent representations, autoencoders or probabilistic models with circadian priors. Harmonic regression, nonparametric tests, count-based models and single-cell approaches identify rhythmic genes and estimate their temporal parameters(9). These methods differ in assumptions, input resolution and prior information, so their outputs need not coincide. Objective evaluation requires known cell phases, rhythmicity labels, amplitudes and gene phases, which experimental transcriptomes rarely provide.

Simulated data can provide such a reference, but existing simulation approaches do not fully meet the requirements for circadian single-cell and spatial transcriptomic analyses. General-purpose simulators such as scMultiSim and scDesign3 reproduce transcriptomic structure, trajectories or spatial organization, but do not explicitly model circadian time-specific cellular and gene-expression changes or provide ground truth for cell phase and gene-level phase and amplitude(10, 11). Conversely, Simphony is a feature- and sample-level data simulator that generates rhythmic expression by controlling amplitude and phase, it cannot explicitly resolve cellular temporal heterogeneity or tissue organization(12). A circadian simulator for single-cell and spatial transcriptomics must therefore incorporate rhythmic variation at both the cell/spot and gene levels, while preserving realistic data structure and explicit temporal parameters.

Here, we introduce scCircaSim, which generates circadian single-cell and spatial transcriptomic data from a single-time-point reference. It models supplied cellular phases, rotates their distribution to target Zeitgeber times and propagates specified anchor gene rhythms through a reference-derived association network. The spatial extension retains anatomical coordinates while incorporating regional phases, neighborhood coupling and correlated phase offsets. Together, these components provide controlled temporal variation with known cell or spot phases and gene-level reference parameters. We evaluate simulation fidelity and use the resulting ground truth to examine phase inference, rhythmicity detection and parameter recovery under defined conditions.(Figure1)

## System and methods

### Reference datasets and simulation framework

scCircaSim is implemented in R and uses reference expression data, cell-type or regional annotations, target sampling times, anchor genes and transcription factors. Single-cell simulation additionally requires reference cell phase estimates. We used liver single-cell data from Tong et al. (2025; GSE288579)(13), with ZT2 cells as the reference for simulation at six time points from ZT2 to ZT22 at 4 h intervals. The dataset contained seven cell types; downstream benchmarks focused on reference 4,857 Hep.CV hepatocytes. A single mouse brain section from OSD-352 provided the spatial reference(14).

Starting from these inputs, scCircaSim advances cell or spot phases to target times and calculates the corresponding changes in anchor gene expression. These changes are propagated through a reference-derived association network and added to reference expression profiles to generate simulated counts. For spatial data, neighborhood coupling and correlated phase offsets modify spot phase progression while tissue coordinates are retained. The outputs include known phases, rhythmicity labels and fitted gene-level parameters for benchmarking.

### Validation and benchmark design

Internal validation comprised three components. First, cell phase fidelity was assessed using circular mean differences, Wasserstein distance, maximum mean discrepancy(MMD) and density overlap. Second, gene expression fidelity was evaluated by anchor amplitude and phase recovery using correlations and error metrics. scRhythmBayes identified rhythmic genes with known simulated phases using FDR < 0.05 and amplitude > 0.25(15), followed by classification, ROC and precision-recall analyses. Finally, parameter sensitivity was examined across gradients of η, controlling rhythmic propagation strength, and λ, controlling ridge regularization in the propagation matrix. These analyses established η = 8 and λ = 0.5 as default settings for subsequent simulations.

Algorithms for circadian cell phase prediction were benchmarked, including TEMPO, CYCLOPS, Cyclum and CoPhaser. TEMPO used a negative binomial model, a 24-point phase grid and anchor gene priors derived from simulated phases; the reference Clock acrophase was used to convert estimates to absolute circadian time(16). An adapted R implementation of CYCLOPS used 30 seed genes, singular value decomposition and the lowest-loss model among 150 fits, calibrated with a 12-gene acrophase panel(17). Cyclum used standardized expression, hidden layers of 30 and 20 units and 1,000 training epochs(18). CoPhaser used 2,000 variable genes plus circadian genes, a 14-gene acrophase prior, a 10-dimensional latent representation and one harmonic, trained for 200 epochs(19). Ground-truth anchor information supported temporal calibration where applicable. Shortest circular errors, empirical cumulative distribution functions (ECDFs), mean absolute error (MAE), root mean squared error (RMSE) and mean cosine agreement quantified recovery of assigned cell phases.

Gene-level benchmarks fixed the period at 24 h. We evaluated three algorithms, using their respective workflows. dryR, HarmonicRegression (HR), and JTK_CYCLE (JTK) were applied to pseudobulk profiles generated by aggregating the simulated single-cell data at each ZT. dryR fitted a single-condition count model after retaining genes with at least 10 counts at two time points(20). HR used log1p-transformed expression normalized to 10,000 counts, with model normalization but no polynomial normalization or trend removal(21). JTK excluded invariant genes(22), used Benjamini–Hochberg-adjusted significance and shifted phase lags by 2 h; raw counts were used for ROC ranking and log-normalized expression for significant-gene calls. Genes were selected at adjusted P < 0.05 and amplitude > 0.25 on each method’s native scale. Detection metrics included genes available to each method, whereas parameter comparisons used significant genes only. ROC ranking used negative log-transformed significance values, with ADJ.P for JTK. Precision, recall and specificity assessed positive-call reliability, rhythmic-gene recovery and non-rhythmic-gene rejection, while accuracy and F1 summarized classification. Mesor and amplitude were evaluated using MAE, median absolute error(MedAE), RMSE and squared Pearson correlation, and phase using circular errors and correlation; JTK was excluded from mesor evaluation.

## Algorithm

### Cell phase generation and anchor rhythms

For each cell type c, cellular phases follow a von Mises distribution:

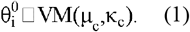

Its center μc is the circular mean of the supplied phases rather than necessarily the sampling ZT, and κc is estimated through the mean-resultant-length relation(23). If cell numbers or composition change, baseline phases are sampled from the fitted distribution and matched to the nearest reference phase within the same type; expression profiles may be reused. Otherwise, supplied phases and profiles are retained.

Progression to target time t follows

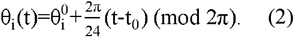

Angles are wrapped to one cycle. Anchor genes expression and changes from baseline are

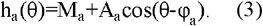

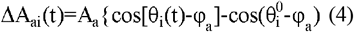

Here, Ma, Aa and φa denote mesor, amplitude and phase. Anchor panels and parameters may be user-defined; amplitudes can be fixed, sampled within a range or assigned individually, and phases may be sampled across 24 h or specified from prior knowledge. These parameters are then used to determine the expression changes of anchor genes across different time points.

### Network construction and expression propagation

Principal component regression (PCR) estimates associations within each cell type and constructs a cell-type-specific reference-derived association network:

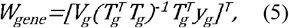

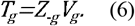

*Z*_*−g*_ contains standardized expression of genes other than target g, *V*_*g*_ contains principal component weights, *T*_*g*_ contains the component scores and *y*_*g*_ denote the target expression. Mapping regression coefficients back through *V*_*g*_ yields the target row of *W*_*gene*_(24, 25). The grouped implementation defaults to ten bootstrap samples of 500 cells and three components per regression. Weights below the 95th percentile in absolute magnitude are removed. Optional canonical polyadic tensor decomposition, with default rank five, extracts shared structure; reconstructed networks are averaged and scaled by their largest absolute entry.

Transcription factor(TF) source columns are retained and each row is normalized by the sum of its absolute weights, giving *W*_*norm*_. Anchor rows form *W*_*A*_, from which the propagation matrix is calculated:

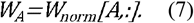

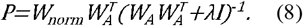

*P*_*ga*_ relates anchor *a* to gene *g* through shared TF associations. Multiplication by the anchor-change matrix converts these relationships into gene expression increments. The ridge term stabilizes inversion(26), with λ = 0.5 by default. Genes are ranked by their total absolute anchor connection:

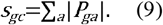

A user-specified fraction ρ_*c*_ receives rhythmic labels *z*_*gc*_ = 1, prioritizing higher scores; other genes receive zero. For example, ρ_*c*_ = 0.6 designates 60% of genes as rhythmic. Propagation then gives

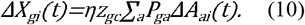

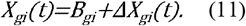

*B*_*gi*_ is matched reference expression. The factor η rescales propagation relative to the PCR-derived weights and defaults to eight. With anchor preservation enabled, directly calculated anchor changes replace propagated anchor increments.

Counts are reconstructed using matched reference library size *L*_*i*_:

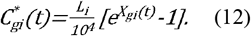

Negative values are set to zero and counts are rounded to integers. Outputs retain source identifiers and phases across cell types and times.

### Gene-level reference parameters

Known simulated phases support harmonic regression on simulated log-normalized expression(21):

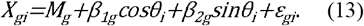

Least squares yields the intercept and harmonic coefficients, defining

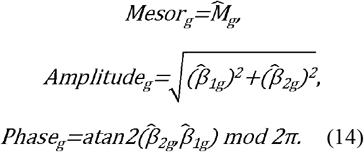

Phase is converted to hours by multiplying by 24/(2π). These fitted characteristics complement the imposed labels and known cell phases for downstream evaluation.

### Spatial phase progression

Regional labels replace cell types for network construction and propagation. Initial spot phases are sampled around the input sampling time:

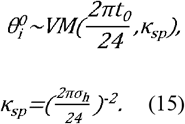

Here, 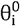 denotes the initial circadian phase of spot i in radians, t_*0*_. is the input sampling time, and κ_sp_ is the concentration parameter controlling the dispersion of spot phases around t_*0*_. The concentration control *σ*_*h*_ defaults to 2 h. Neighborhood weights may use coordinates, coordinates combined with BANKSY representations, or a supplied matrix, and are symmetrized before use. Coordinate-based weights are

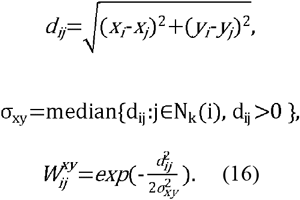

Here, *d*_*ij*_ is Euclidean distance and *N*_*k*_(i) contains the k nearest spots, with k = 6 by default. The median positive local distance sets the spatial bandwidth, limiting sensitivity to unusually large gaps. Nearby spots therefore receive stronger coordinate-based weights. BANKSY-based weights additionally incorporate representation similarity(27):

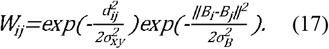

The bandwidth *σ*_*B*_ is the median positive representation distance over the same neighbor pairs; BANKSY’s mixing parameter defaults to 0.2. Target phases incorporate neighborhood coupling and correlated offsets:

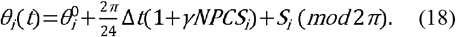

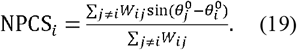

Δt = (t − *t*_*0*_) mod 24 and *γ* defaults to 0.2. Neighborhood phase coupling score (NPCS) is calculated once from initial phases, excluding self-weights, and is zero without positive neighbor weight. For non-negative weights, NPCS_*i*_ ranges from −1 to 1, with its sign determining whether local phase progression is accelerated or decelerated, while *γ* controls the strength of this neighborhood-dependent modulation.

Correlated offsets S follow

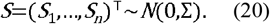

*Σ* is obtained by projecting symmetric W to a positive-definite correlation matrix with unit diagonal. Correlated offsets allow strongly connected spots to vary together, while the zero mean avoids a systematic phase shift across repeated simulations. One Gaussian field is shared across non-reference target times; reference phases remain unchanged. The resulting phases determine anchor changes, followed by the same propagation and count reconstruction as in single-cell simulation.

## Implementation

### Preservation of phase distributions and rhythmic expression

scCircaSim supports the simulation of multi-time-point single-cell transcriptomic data from a single-time-point reference. After cell-phase estimation, cells phases are rotated to user-defined target times, and anchor expression alterations are propagated through the PCR network to simulate expression at one or multiple time points (Figure 2A). A liver single-cell dataset with six time points was used to evaluate scCircaSim, with ZT2 selected as the single-time-point reference for generating a multi-time-point circadian dataset. The simulated liver data retained the overall quality characteristics of the reference dataset (Figure S1A-D). Anchor and propagated rhythmic genes oscillated over the day, whereas non-rhythmic genes remained comparatively stable (Figures S1E,F and S2A)

**Figure 1.**
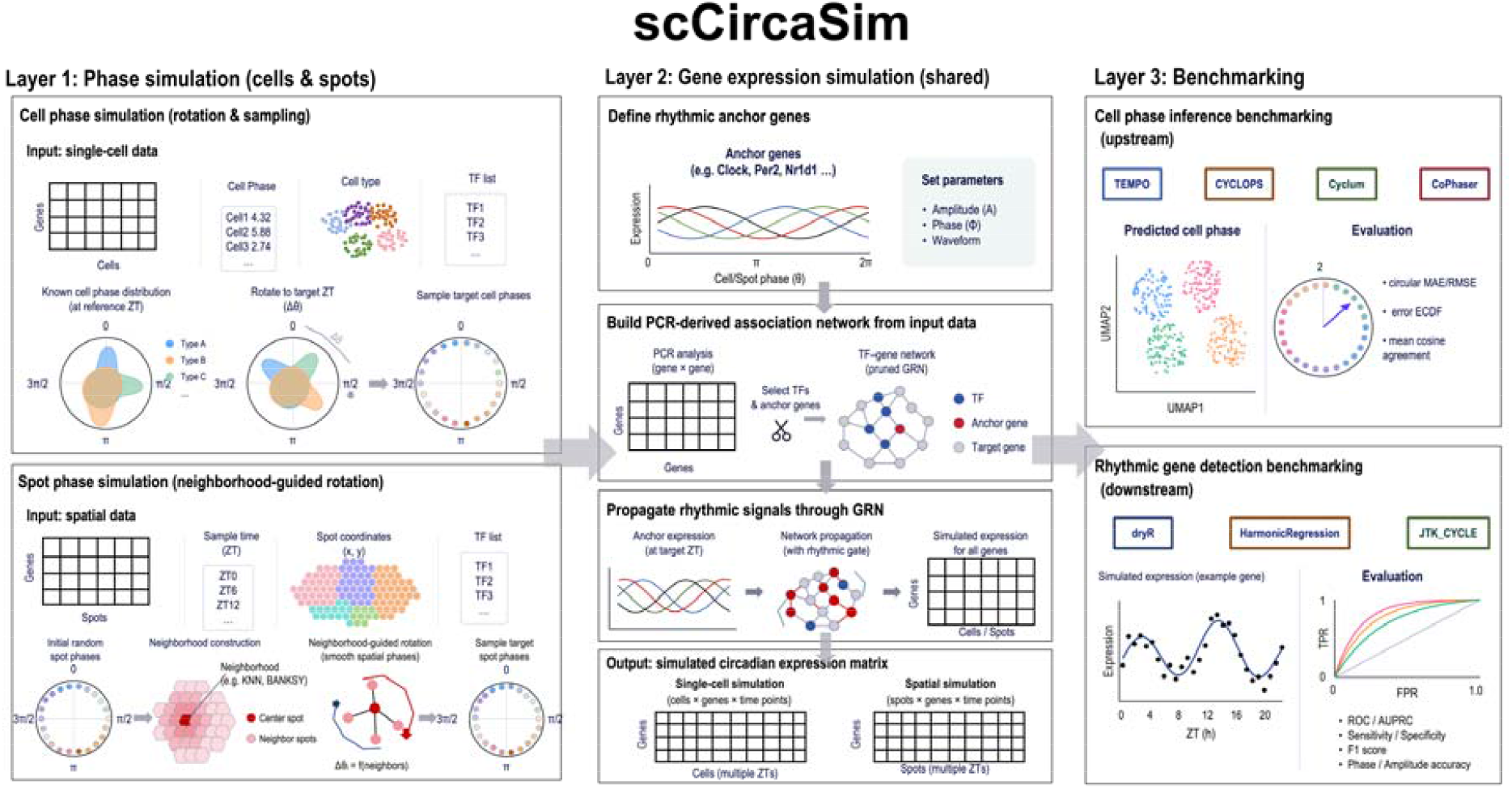
Overview of circadian simulation and benchmarking with scCircaSim. The framework combines cell or spot phase simulation, shared gene expression simulation and downstream benchmarking. Cellular phases are rotated to target times; spatial phases additionally incorporate neighborhood coupling and correlated offsets. Anchor rhythms are propagated through a PCR-derived association network to generate expression profiles with known temporal characteristics. Simulated phases and gene-level references support evaluation of phase inference, rhythmicity detection and parameter recovery.

**Figure 2.**
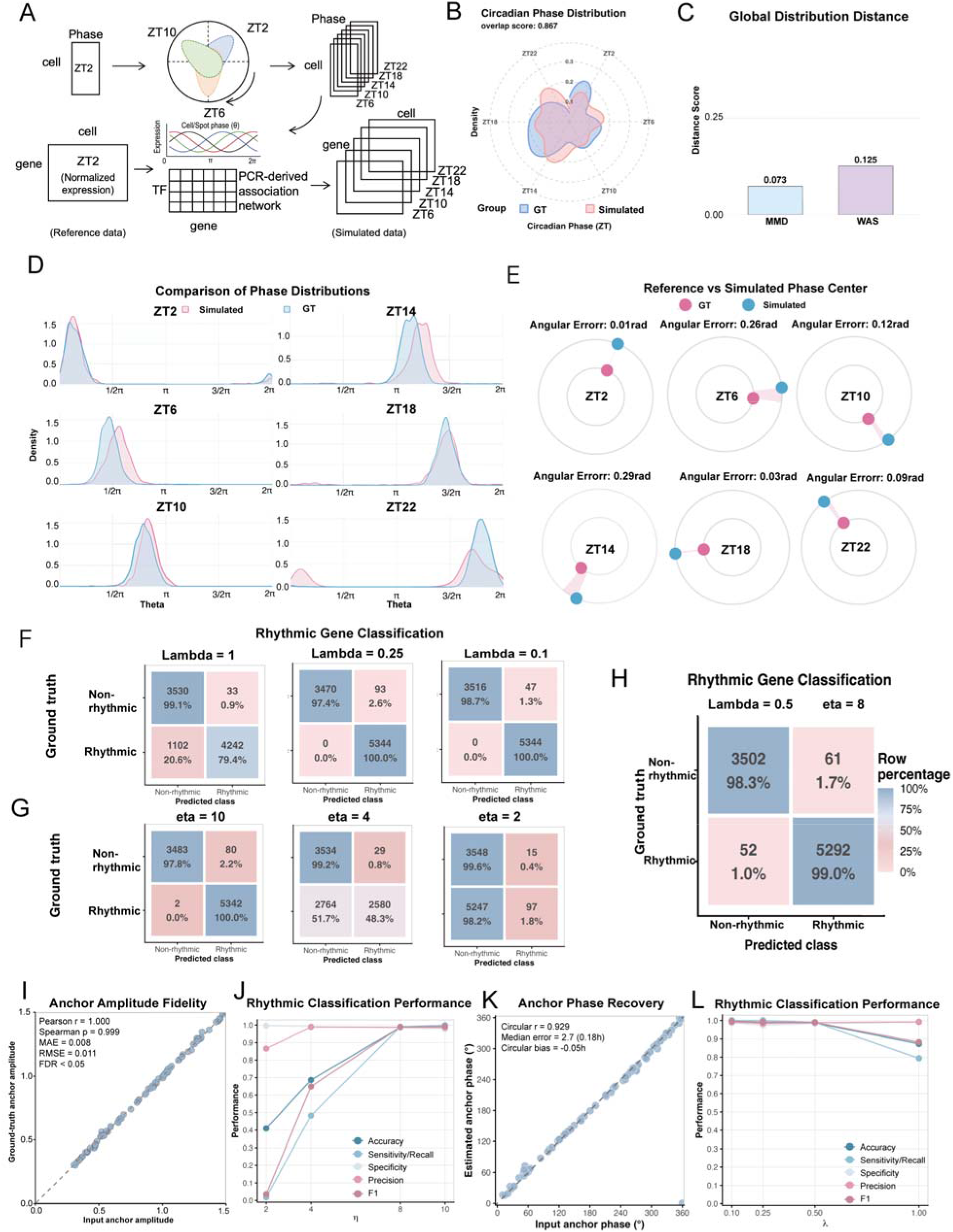
Validation of phase progression and rhythmic expression in simulated single-cell data. (A)Workflow for generating simulated single-cell transcriptomes at multiple time points from a single-time-point reference. (B)Pooled reference and simulated phase densities and overlap. (C) Global MMD and Wasserstein distance. (D) Phase densities at each ZT. (E) Circular mean phases and angular differences. (F, G) Confusion matrices for rhythmic gene classification at different regularization parameters λ(F)and propagation strengths η(G), with counts and percentages within each ground-truth class. (H) Inferred rhythmicity labels versus imposed ground truth. (I) Assigned versus fitted anchor amplitudes. (J) Classification metrics across propagation strengths η. (K) Assigned versus estimated phases among detected anchors. (L) Classification metrics across regularization parameters λ.

To validate cell-phase recovery after rotation, pooled simulated and reference phase distributions showed an overlap of 0.867, MMD of 0.073 and Wasserstein distance of 0.125 (Figure 2B,C). Across individual time points, Wasserstein distances were below 0.3, with the smallest values at ZT10 and ZT18 except for the resampled ZT2 reference (Figure S2B). MMD showed a similar pattern with relatively low distances across time points (Figure S2C). At each sampling time, simulated and reference phase distributions showed substantial overlap(Figure 2D). The largest circular mean difference was 0.29 rad (approximately 1.11 h) at ZT14, followed by 0.26 rad at ZT6 and 0.12 rad at ZT10; differences at the remaining times were below 0.10 rad. (Figure 2E). Simulated cell-type proportions matched the reference population (Figure S2D), while agreement with upstream phase estimates further supported retention of the reference phase structure (Figures S2E,F).

For gene expression, PCR-based propagation of anchor rhythms was evaluated across gradients of η and λ to determine default parameters. λ had little effect on the number of rhythmic genes, but the true-positive rate declined sharply at λ = 1 (Figure 2F). Increasing η strengthened rhythmic expression and increased significant rhythmic genes, whereas excessive propagation increased false negatives (Figure 2G). At λ = 0.5 and η = 8, 99.0% of designated rhythmic genes and 98.3% of non-rhythmic genes were correctly classified, with anchor amplitudes preserved (Figure 2H,I). Median anchor amplitude error was approximately 0.18 h (Figure 2K,S2F). Classification metrics plateaued at η = 8 (Figures 2J, S2G), while stronger regularization reduced sensitivity despite high precision (Figure 2L) and decreased AUROC (Figure S2H). We therefore selected λ = 0.5 and η = 8 as default parameters. Together, these results support controllable rhythmic simulation while retaining the reference phase distribution.

### Comparative evaluation of cell phase inference

Four phase inference workflows were evaluated in Hep.CV hepatocytes against the assigned phases. TEMPO positioned distributions nearer the target ZTs; Cyclum estimates were more dispersed, CYCLOPS covered a narrower phase range, and CoPhaser retained differences after anchor-based alignment (Figure 3A,B). At the individual-cell level, TEMPO estimates broadly followed the diagonal with appreciable dispersion. CoPhaser showed a largely reversed phase trajectory and CYCLOPS concentrated estimates in restricted intervals under the evaluated calibration. TEMPO most closely reproduced the pooled phase density, illustrating that distributional agreement and individual-cell accuracy capture different properties (Figure 3C). For individual cells, normalized error-ECDF area was 0.786 for TEMPO, compared with values near 0.5 for the other workflows (Figure 3D). Typical errors were concentrated near 2 h for TEMPO, versus approximately 6 h for the remaining methods (Figure 3E). MAE was 2.57 h for TEMPO, and exceeded 5 h for CoPhaser, Cyclum and CYCLOPS; RMSE gave the same ordering (Figure 3F,G). Mean cosine agreement was 0.70 for TEMPO, and near zero for the other methods (Figure 3H). Thus, TEMPO showed closer absolute phase recovery in this simulated setting, with interpretation conditional on each workflow’s temporal inputs and calibration.

**Figure 3.**
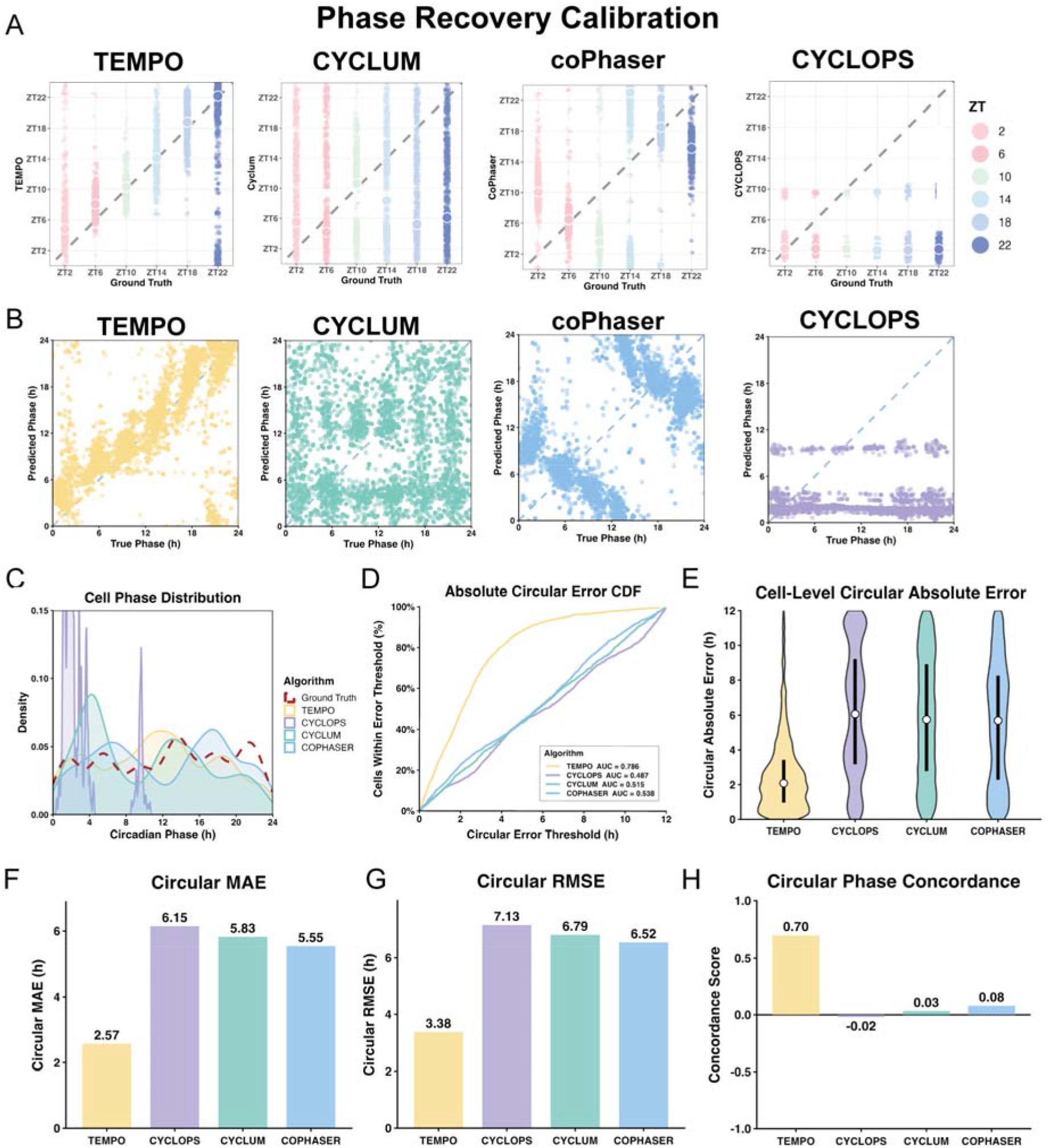
Benchmarking cell phase inference against simulated ground truth. (A) Phase estimates grouped by target ZT for TEMPO, Cyclum, CoPhaser and CYCLOPS, from left to right. (B) Inferred versus simulated phases for TEMPO, Cyclum, CoPhaser and CYCLOPS; diagonals indicate agreement. (C) Inferred and ground-truth phase densities. (D) Cumulative absolute circular error distributions; legends report areas normalized over 0 to 12 h. (E) Cell-level absolute circular errors. (F) Circular MAE. (G) Circular RMSE. (H) Mean cosine of phase differences.

### Rhythmicity detection and parameter recovery

The simulated gene-level ground truth enabled benchmarking of rhythmic gene detection and parameter estimation. With 60% of genes designated rhythmic, dryR, HR and JTK identified 91.5%, 47.4% and 2.2% as rhythmic, respectively (Figure 4A). dryR extensively overlapped with HR, whereas the smaller JTK set largely overlapped with both methods (Figure 4B). Ground-truth classification was used for further evaluation. ROC discrimination was high for HR, JTK, with JTK highest, but lower for dryR (Figure 4C). At the selected thresholds, dryR recovered 87.1% of rhythmic genes but misclassified 3,455 non-rhythmic genes; HR achieved 75.6% sensitivity and 95.0% specificity, whereas JTK showed high specificity but limited sensitivity (Figure 4D). Parameter recovery was assessed within each method’s detected set. HR showed lower mesor errors than dryR, with mesor R^2^ of 0.940 for HR and 0.783 for dryR (Figure 4E). Amplitude MAE was 0.123 for JTK, 0.332 for HR and 1.491 for dryR, with similar MedAE and RMSE rankings (Figure 4F). JTK’s favorable agreement, however, applies to its small detected subset, and native parameter scales affect cross-method comparisons. For gene phase, circular MAE and RMSE were lower for HR. dryR had lower MedAE than JTK despite higher MAE, indicating different error distributions. JTK showed stronger circular correlations than HR and dryR (Figure 4G). Accordingly, relative phase association need not imply equally close absolute phase recovery. The common reference supports joint evaluation of detection and parameter estimation without reducing performance to a single ranking.

**Figure 4.**
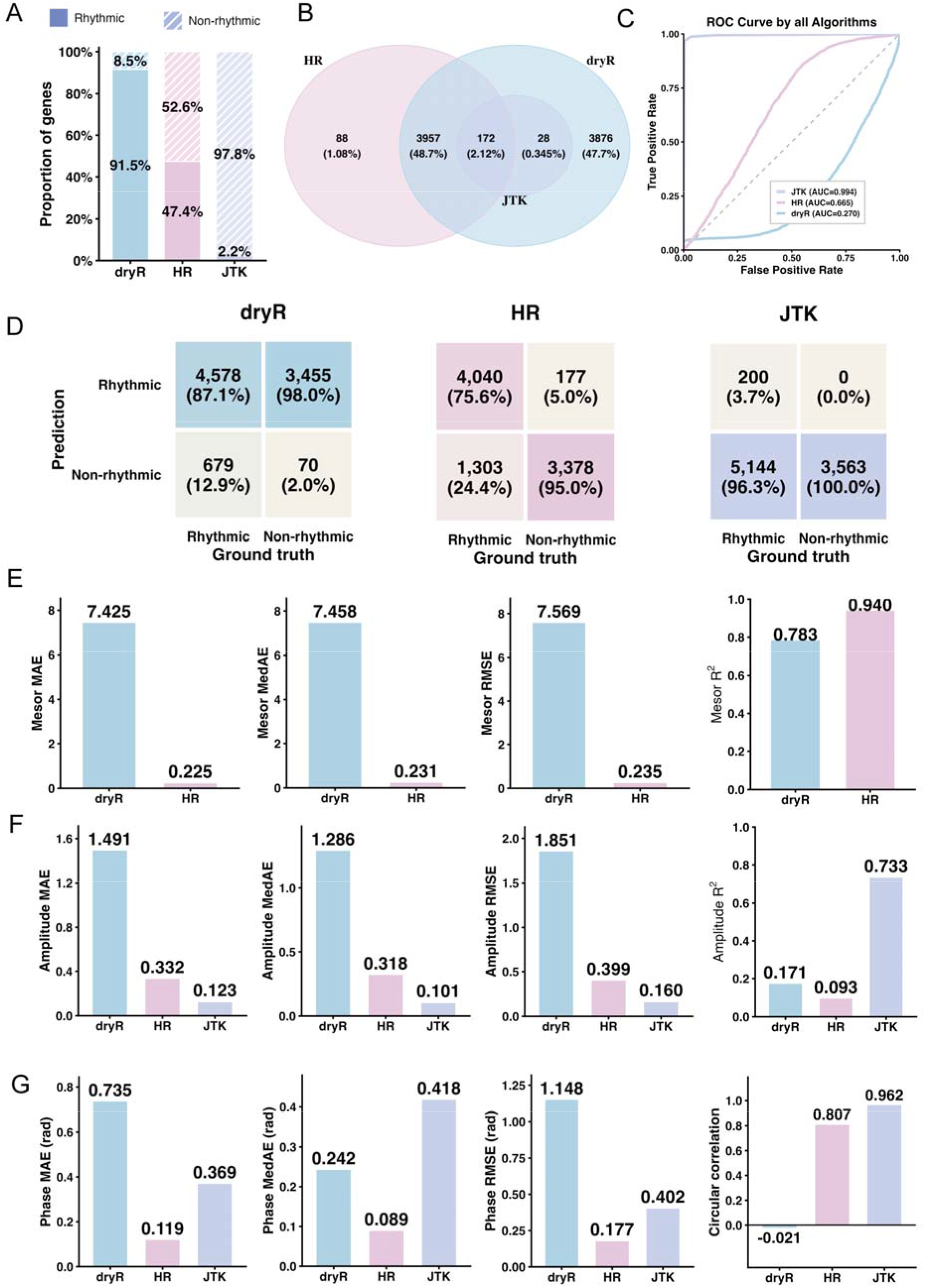
Benchmarking rhythmicity detection and parameter recovery. (A) Rhythmic fractions called by dryR, HarmonicRegression (HR) and JTK_CYCLE (JTK). (B) Overlap of detected sets. (C) ROC curves and AUCs. (D) Confusion matrices, with counts and percentages within ground-truth classes. (E, F) Mesor and amplitude agreement summarized by MAE, MedAE, squared Pearson correlation and RMSE; JTK is excluded from mesor evaluation. (G) Circular phase MAE, MedAE, correlation and RMSE, with errors in radians. Parameter comparisons use each method’s detected genes and native scales.

### Spatial simulation across brain regions

A single annotated mouse brain section supported simulation at six time points while retaining anatomical organization (Figure 5A,B). Quality metrics and embeddings are shown in Figure S3A-J. Regional initial phases followed von Mises distributions (Figure S3K), and simulated densities remained concentrated near target sampling times (Figures 5C and S4A). NPCS summarized spot-level neighborhood phase adjustments within regions. Most regional distributions were approximately symmetric, with shifts in some regions consistent with local advancement or delay (Figure S4B). The correlated spatial effect showed no significant departure from normality in the Shapiro-Wilk test (P = 0.413; Figure 5D). Pooled and time-specific expression distributions remained broadly similar to the reference (Figures 5E and S4C).

**Figure 5.**
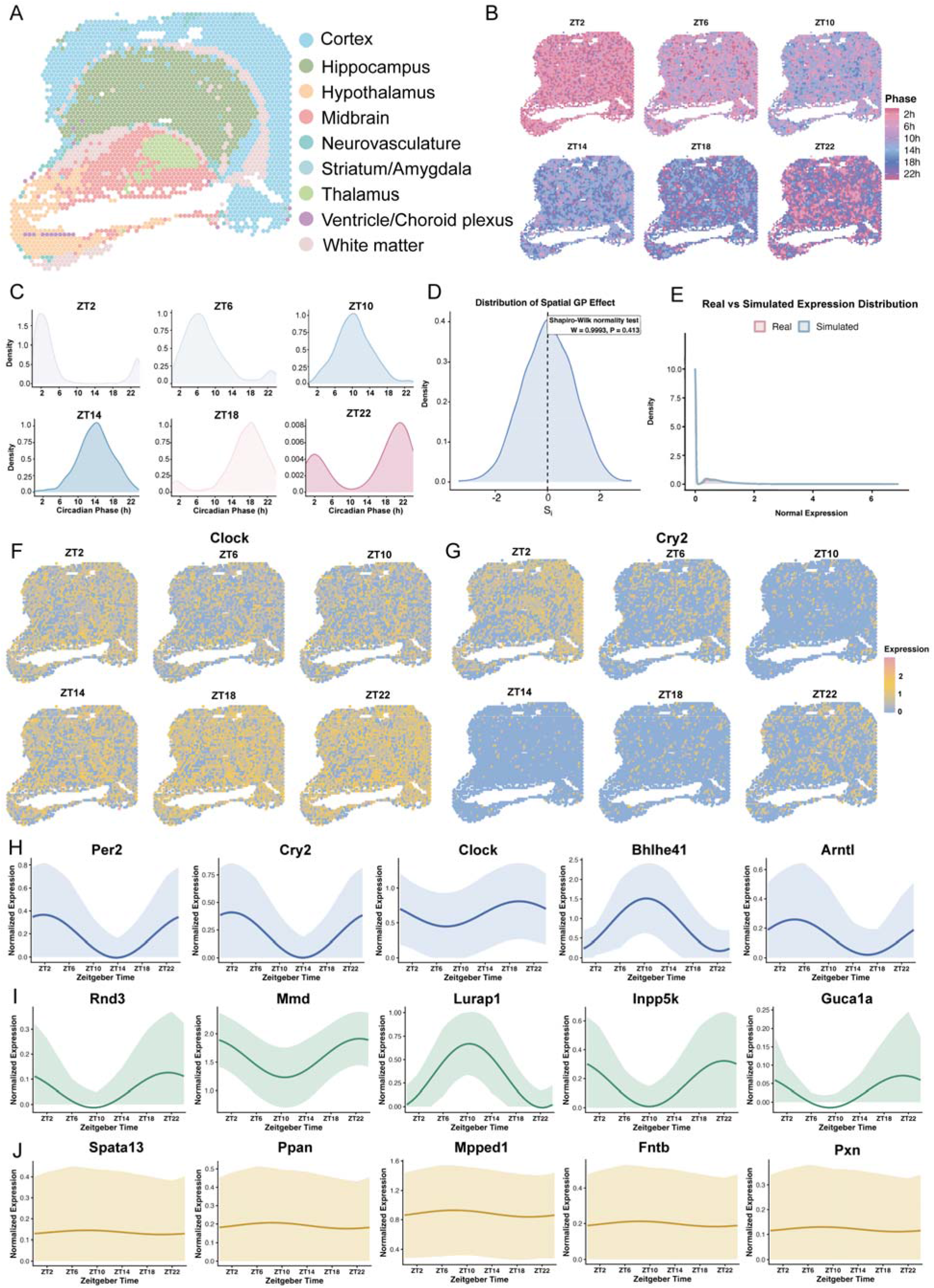
Circadian phase and expression simulation from one brain section. (A) Reference section colored by region. (B) Simulated phases at ZT2, ZT6, ZT10, ZT14, ZT18 and ZT22. (C) Corresponding phase densities. (D) Simulated spatial Gaussian effect; Shapiro-Wilk W = 0.9993, P = 0.413. (E) Reference and pooled simulated expression distributions. (F, G) Spatial expression of Clock and Cry2 across simulated times. (H-J) Temporal profiles of anchor, rhythmic non-anchor and non-rhythmic genes. Curves show harmonic fits to mean normalized expression; bands show fitted curves plus or minus interpolated standard deviations across spots.

Simulated circadian gene expression varied across anatomical regions. Clock expression was lower at ZT6 and ZT10 and increased between ZT14 and ZT22; Cry2 was higher at ZT2 and ZT22 and lower during the intervening portion of the cycle, with regional variation (Figures 5F,G and S4D). Local variation persisted within regions (Figure S5A). Anchor and propagated rhythmic genes showed temporal oscillations, whereas non-rhythmic genes remained relatively stable (Figures 5H-J and S5B-D). These patterns illustrate generation of specified circadian expression states within a common spatial reference, with both regional organization and gene-specific temporal behavior represented.

## Discussion

scCircaSim connects controlled circadian simulation with quantitative benchmarking. Its central requirement is to represent periodicity across cells, genes and spatial neighborhoods while retaining an identifiable temporal reference.

Nevertheless, one snapshot cannot uniquely identify a complete circadian trajectory or establish whether expression will subsequently rise or fall(28). Simulated time points therefore represent specified temporal scenarios. Random and user-defined parameters allow exploration informed by prior knowledge, while known labels and fitted parameters make those scenarios testable. Harmonic parameterization describes 24 h rhythms tractably; asymmetric waveforms, transient responses and multiple periodic components would require additional assumptions.

However, gene expression programs do not operate independently, and assigning amplitude and phase to each gene separately would neglect the gene–gene dependencies present in the reference transcriptome. Thus, scCircaSim provides a set of established rhythmic anchor genes(29), while also allowing users to define custom anchors. These anchor genes provide predefined rhythmic trajectories with controllable temporal parameters, including amplitude and phase, whereas the PCR-derived network determines how strongly these temporal signals are transmitted to other genes. Temporal changes in anchor gene expression are propagated to non-anchor genes according to their relationships in the reference-derived PCR network, such that changes in anchor expression across circadian phases produce coordinated changes in downstream genes. This anchor-to-network strategy therefore links rhythmicity in non-anchor genes to the temporal variation of anchor genes rather than assigning independent oscillations to each gene, while retaining gene dependencies derived from the reference transcriptome. Propagation parameters control its detectability: increasing η improved recovery before a plateau, whereas λ = 1 attenuated sensitivity relative to moderate regularization. Because effective propagation depends on PCR weights, normalization and regularization, η should be chosen relative to the reference network rather than interpreted as a universal biological constant.

scCircaSim first normalizes the reference data to reduce differences in sequencing depth across cells, allowing rhythmic perturbations to represent relative expression changes rather than variation driven by library size. Rhythmic perturbations are then introduced in the normalized expression space, and the resulting expression profiles are converted back to count-like values using the library sizes of the matched reference cells. Because the magnitudes of negative rhythmic deviations can exceed the baseline expression of lowly expressed genes, reconstructed values may fall below the biological lower bound of zero. These values are therefore truncated at zero before integer rounding, introducing additional zeros and potentially compressing the troughs of simulated rhythms, particularly for genes with low baseline expression or large amplitudes. The reconstructed count matrix is subsequently re-normalized, from which mesor, amplitude and phase are fitted to characterize the realized temporal patterns and serve as ground truth for downstream evaluation.

The phase benchmark also distinguishes learning circular structure from calibrating it to biological time. TEMPO uses probabilistic inference and clock priors; CYCLOPS reconstructs an eigengene-based cycle; Cyclum learns circular pseudotime; CoPhaser here separates periodic and contextual variation. Ground-truth anchors provided priors or calibration where supported, including conversion of TEMPO phases to absolute time. This reduces coordinate ambiguity without making the workflows’ information inputs equivalent. Sampling time itself also differs from cell-specific phase. TEMPO performed favorably with clock priors. The larger absolute errors of the remaining workflows characterize their behavior under this design, without excluding their utility for other periodic structures or datasets. In particular, an unsupervised cycle need not correspond directly to circadian time, and anchor calibration alone does not guarantee recovery of cell-level variation.

Gene-level benchmarking also revealed method-specific trade-offs. Although pseudobulk aggregation can attenuate rhythms arising from cellular phase heterogeneity, it does not fully explain differences among methods applied to the same data(30). JTK favored specificity, HR showed more balanced detection, whereas dryR identified more rhythmic genes but with more false positives. Parameter recovery further showed that detection performance alone does not capture all aspects of method accuracy, highlighting the need to jointly consider rhythmic gene discovery and the recovery of mesor, amplitude and phase. For phase recovery, circular correlation additionally captures the preservation of relative phase relationships, which may remain consistent despite systematic offsets that increase absolute error(31). Overall, the benchmark showed distinct performance profiles across JTK, HR and dryR, with no single method consistently outperforming the others across rhythmic gene detection and circadian parameter recovery. By providing predefined rhythmicity labels and corresponding circadian parameters, scCircaSim enables these complementary aspects to be evaluated against a common ground truth, thereby facilitating characterization of method-specific performance under controlled circadian conditions.

Spatial simulation addresses a further practical constraint because profiling different time points usually requires different sections or specimens, inevitably introducing inter-sample variation(32). Generating temporal states on a fixed coordinate map supports controlled comparisons with anatomy held constant, although these remain conditional simulations rather than longitudinal measurements. Assigning a common phase to all spots would fail to represent local temporal heterogeneity, whereas independent phase sampling would discard neighborhood relationships. In scCircaSim, regional phase distributions retain local heterogeneity, neighborhood coupling represents local adjustments, and correlated offsets introduce shared spatial variation. NPCS specifically describes net phase advancement or delay, rather than a general measure of synchronization. Together, these components support exploration of regional temporal programs and spatial phase relationships within a common tissue reference.

Thus, scCircaSim supports reproducible benchmarking and preliminary study design. Configurable trajectories, sampling schedules and analytical choices can help prioritize informative experiments, reduce avoidable resource use and potentially limit unnecessary animal experiments. These benefits depend on treating simulated dynamics as hypotheses for validation and evaluating conclusions within the reference data and assumptions used.

## Supporting information

FigureSupplement

## Funding

This work was supported by the Qingdao Key Technology Research and Industrialization Demonstration Program (26-1-1-gjgg-66-nsh). We thank our laboratory members for helpful discussions and technical support, the researchers who generated the public datasets, and the developers of the benchmarked computational tools.

## Data availability

Single-cell RNA sequencing data are available from GEO under GSE288579. Brain spatial data are available from NASA GeneLab within the Open Science Data Repository under OSD-352, with processed spatial count matrices from Mendeley Data. No new sequencing data were generated.

## Code availability

scCircaSim is implemented in R and available at https://github.com/Liwei-DynamicsLab/scCircaSim.

Third-party software is available from its respective developer.

## Notes

https://github.com/Liwei-DynamicsLab/scCircaSim

### Competing Interest Statement

The authors have declared no competing interest.

