## Supplementary material for "scCircaSim: A Circadian Simulator for Single-Cell and Spatial Transcriptomics": FigureSupplement

### Supplementary Figure S1

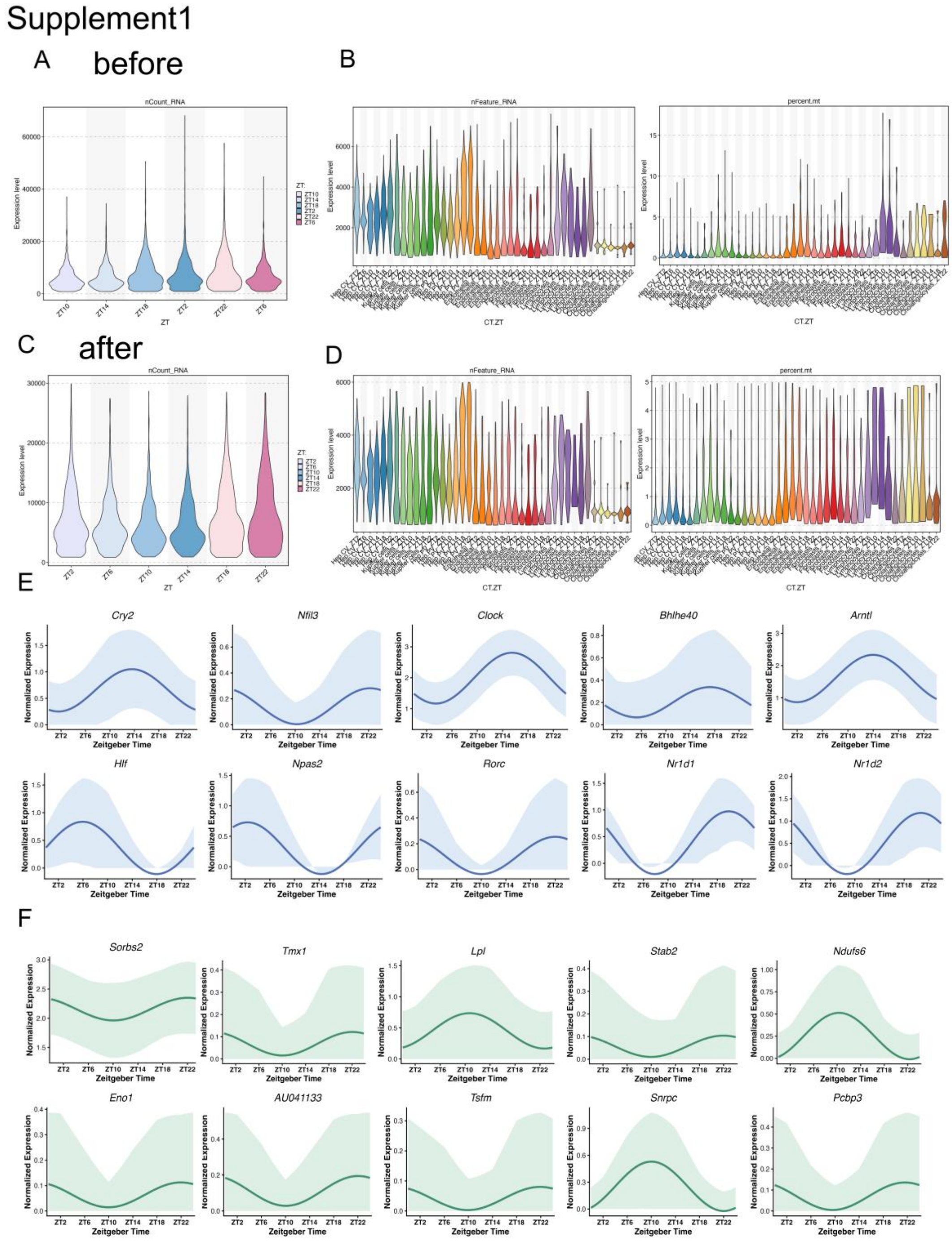


**Supplementary Figure S1 Quality assessment and rhythmic expression profiles in simulated single-cell data**

(A, B) Quality metrics before filtering: total RNA counts per cell (nCount_RNA) by ZT (A), and numbers of detected genes (nFeature_RNA) and percentages of mitochondrial counts (percent.mt) by cell type and ZT (B). (C, D) Corresponding metrics after filtering. Violin widths represent the density of cells at each value. ZT, Zeitgeber time. (E-F) Representative normalized expression profiles of rhythmic anchor genes (E) and rhythmic non-anchor genes (F) over a 24 h cycle.

### Supplementary Figure S2

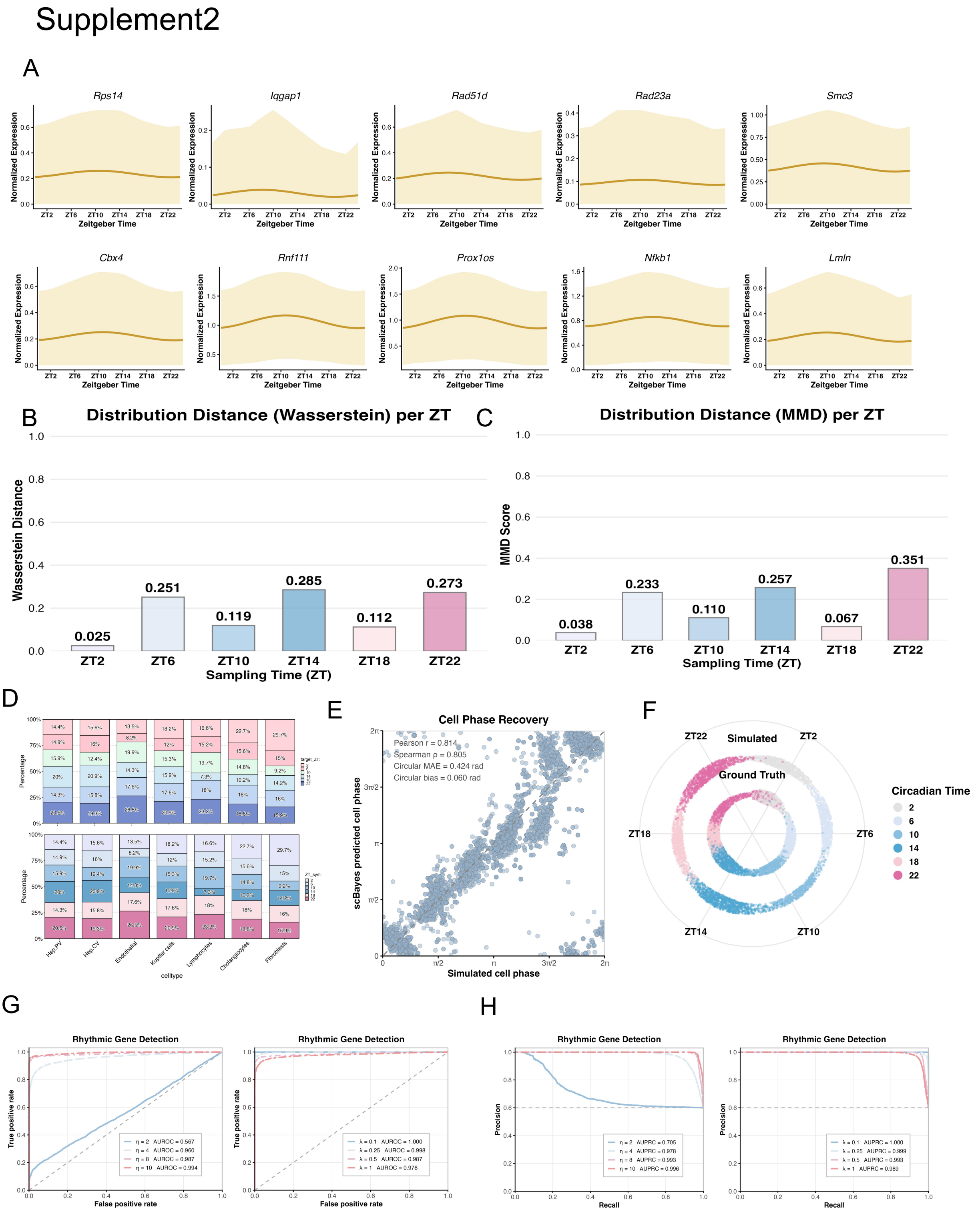


**Supplementary Figure S2 Nonrhythmic expression profiles and validation of cell phases and propagation parameters** (A) Representative normalized expression profiles of non-rhythmic genes over a 24 h cycle. Gene names are shown above individual panels. Solid curves represent harmonic fits, and shaded bands indicate the fitted curves plus or minus the standard deviation across cells, interpolated between sampling times and truncated at zero at the lower bound. ZT, Zeitgeber time. (B,C)

Wasserstein distance and MMD at each ZT. (D) Cell-type proportions. (E) Simulated phases versus scRhythmBayes estimates. (F) Circular scatter plot of reference and simulated cell phases, colored by ZT. (G) ROC curves for the η and λ sensitivity analyses, with areas under the ROC curves (AUROCs) indicated. (H) Corresponding precision-recall curves, with areas under the precision-recall curves (AUPRCs) indicated.

**Supplementary Figure S3**

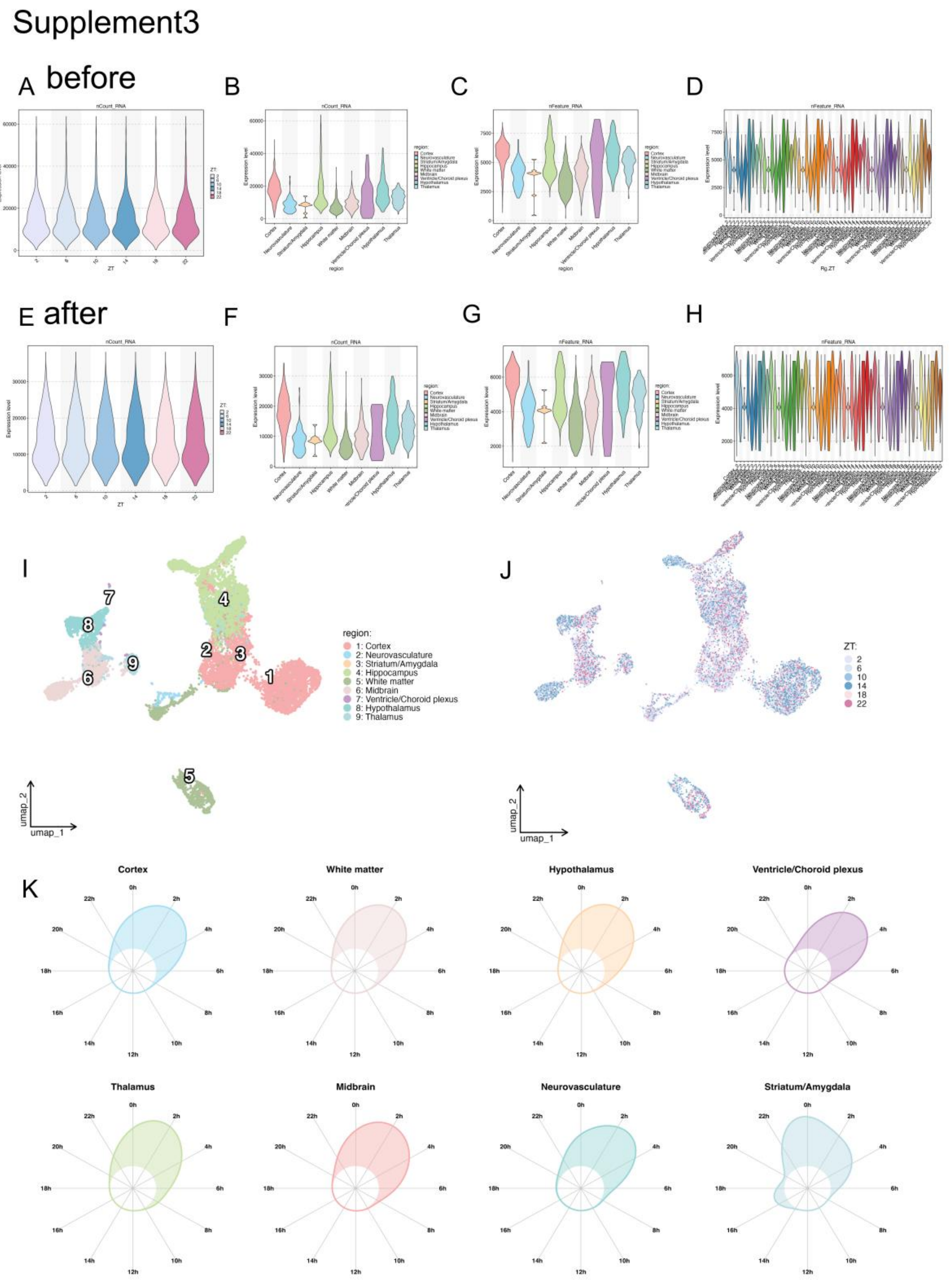


**Supplementary Figure S3 Quality assessment and regional structure of simulated spatial transcriptomic data**

(A-D) Quality metrics before filtering: total RNA counts per spot by ZT (A) and anatomical region (B), and numbers of detected genes per spot by region (C) and by region and ZT (D). (E-H) Corresponding metrics after filtering. Violin widths represent the density of spots at each value. (I, J) UMAP of simulated spots pooled across time points, colored by anatomical region (I) or ZT (J). (K) Polar density plots of initial spot phases in the indicated anatomical regions. nCount_RNA, total RNA counts per spot; nFeature_RNA, number of detected genes per spot; ZT, Zeitgeber time.

**Supplementary Figure S4**

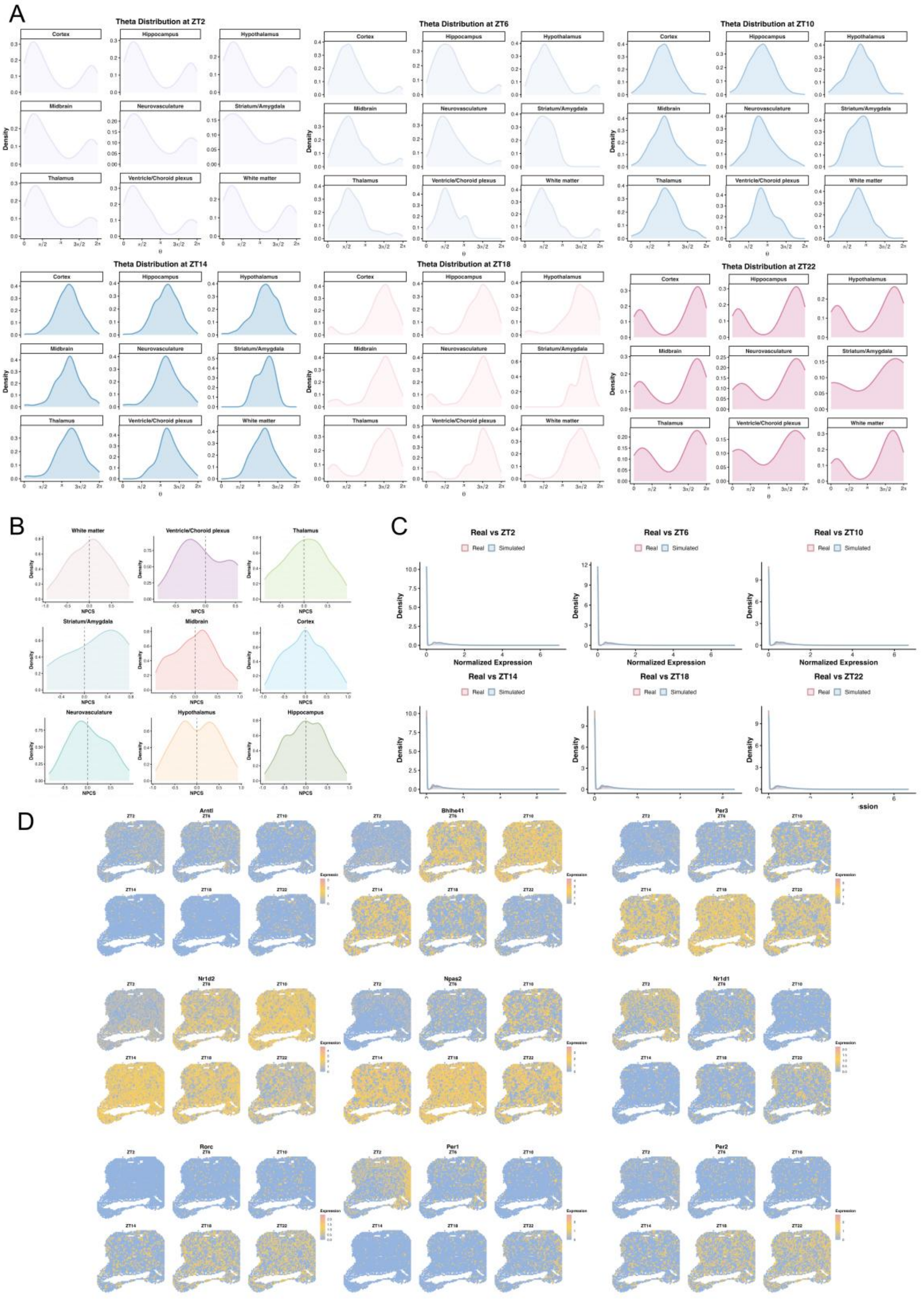


**Supplementary Figure S4 Regional phase distributions and spatial expression patterns in simulated brain data**

(A) Spot phase densities stratified by anatomical region and simulated ZT, with phases expressed in radians. ZT, Zeitgeber time. (B) Distributions of the neighborhood phase coupling score (NPCS) within anatomical regions. (C) Distributions of normalized expression in the reference section and simulated data at each ZT. (D) Spatial expression maps of selected circadian genes across the whole section at six simulated time points. Genes shown are Arntl, Bhlhe41, Per3, Nr1d2, Npas2, Nr1d1, Rorc, Per1 and Per2. Colors indicate normalized expression according to the gene-specific scales. ZT, Zeitgeber time.

**Supplementary Figure S5**

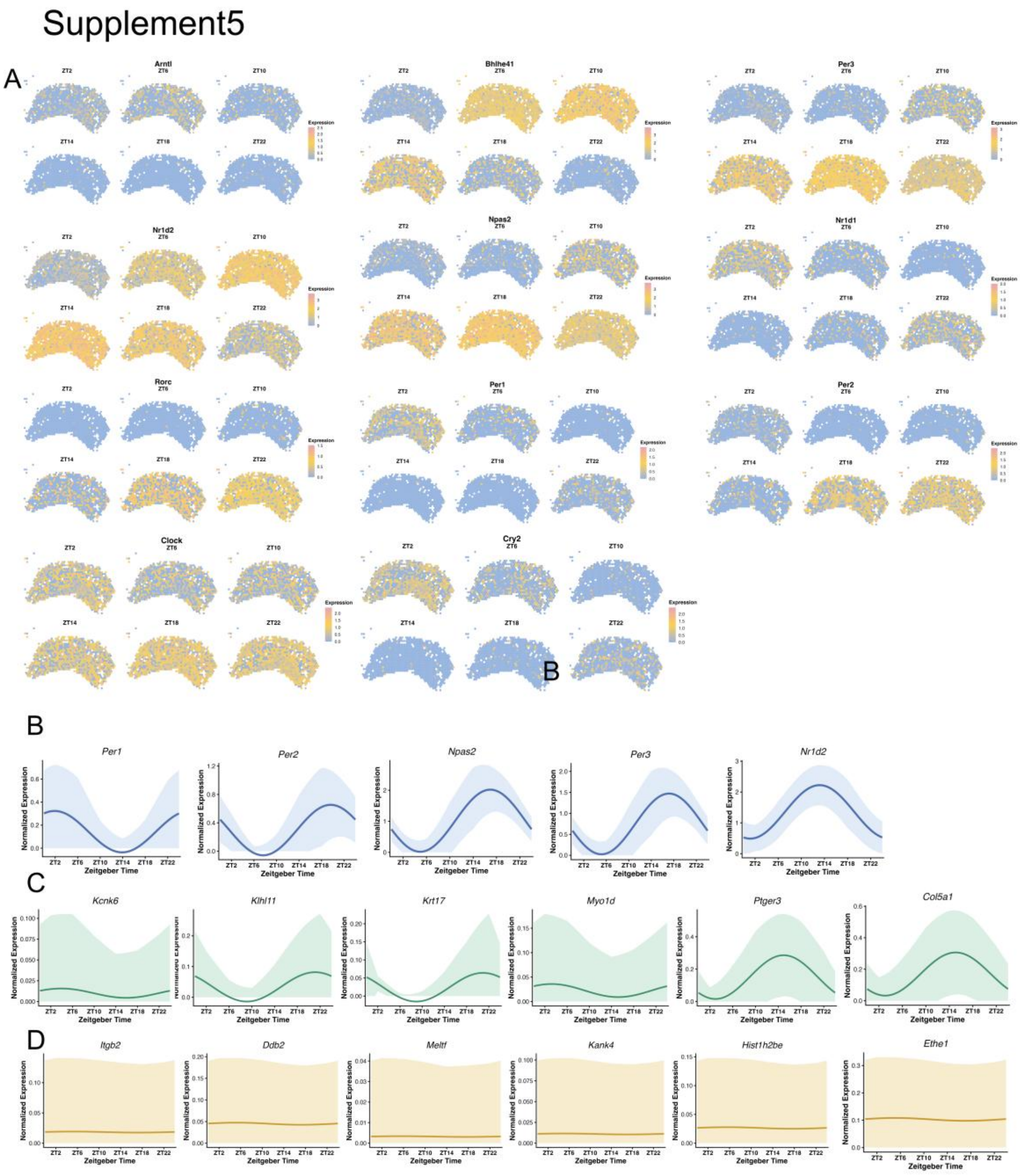


**Supplementary Figure S5 Hippocampal expression maps and temporal profiles of simulated gene expression**

(A) Spatial expression of anchor genes within the hippocampus at six simulated ZT points. (B-D) Temporal expression profiles of additional rhythmic anchor genes (B), rhythmic non-anchor genes (C) and non-rhythmic genes (D) in the simulated spatial data. Solid curves show harmonic fits to mean normalized expression, and shaded bands show the fitted curves plus or minus the interpolated standard deviation across spots, with lower bounds truncated at zero.
